# Computationally reconstructed interactome of *Bradyrhizobium diazoefficiens* USDA110 reveals novel functional modules and protein hubs for symbiotic nitrogen fixation

**DOI:** 10.1101/2021.03.06.434201

**Authors:** Jun-Xiao Ma, Yi Yang, Guang Li, Bin-Guang Ma

## Abstract

Symbiotic nitrogen fixation is an important part of the nitrogen biogeochemical cycles and the main nitrogen source of the biosphere. As a classical model system for symbiotic nitrogen fixation, rhizobium-legume systems have been studied elaborately for decades. Detailed panorama about the molecular mechanism of the communication and coordination between rhizobia and host plants is becoming clearer. For more systematic insights, there is an increasing demand on new studies integrating multi-omics information. Here we present a comprehensive computational framework, integrating the reconstructed protein interactome of *B. diazoefficiens* USDA110 with its transcriptome and proteome data, to study the complex protein-protein interaction (PPI) network involved in the symbiosis system. We reconstructed the interactome of *B. diazoefficiens* USDA110 by computational approaches. Based on the comparison of interactomes between *B. diazoefficiens* USDA110 and other rhizobia, we inferred that the slow growth of *B. diazoefficiens* USDA110 may owe to the requirement of more protein modifications and further identified 36 conserved functional PPI modules. Integrated with transcriptome and proteome data, interactomes representing free-living cell and symbiotic nitrogen-fixing (SNF) bacteroid were obtained. Based on the SNF interactome, a core-sub-PPI-network for symbiotic nitrogen fixation was determined and 9 novel functional modules and 11 key protein hubs playing key roles for symbiosis were identified. The reconstructed interactome of *B. diazoefficiens* USDA110 may serve as a valuable reference for studying the mechanism underlying the SNF system of rhizobia and legumes.

## 1. Introduction

In most cases, rhizobia are a group of gram-negative soil bacteria within the family Rhizobiaceae. They can colonize roots of legumes, establish symbiotic relationship with them and perform nitrogen fixation. This kind of symbiotic nitrogen-fixing (SNF) system can convert inorganic nitrogen to organic nitrogen and constitute an important part of the biogeochemical cycle^1^. As the type strain of Bradyrhizobium, an important genus of rhizobia, *B. diazoefficiens* USDA110 (formerly known as *B. japonicum* USDA110) ^2^ can establish a symbiotic relationship with *Glycine max* (soybean). The *G. max* – *B. diazoefficiens* system has been an important model for the study of SNF system^3,4^. In addition, given the efficient SNF ability of *B. diazoefficiens* USDA110^5^, it is widely used in agriculture and environmental engineering^6,7^. Owing to its importance, various biological characteristics of *B. diazoefficiens* have been extensively studied for decades, especially its SNF mechanism such as the information exchange between *B. diazoefficiens* and *G. max* during the process of establishing symbiosis^8,9^. The genome of *B. diazoefficiens* reference strain USDA110 was completely sequenced in 2002^10^ and corresponding high-throughput omics studies were performed in recent years^11–17^. Although a large amount of knowledge has been obtained through these studies, contribution of the protein-protein interaction (PPI) network of this species in SNF mechanism remains elusive.

The proteins in cells do not exist in isolation. Biological processes often involve multiple proteins and their interactions^18,19^. Thousands of proteins are coordinated, forming a functional network, the PPI network, which is becoming one of the main research objects in systems biology. PPI networks have become effective tools for understanding the cellular behaviors and solving various biological problems in transcription, translation, metabolism, gene regulation and signal transduction^20^. From the more comprehensive perspective of a PPI network, new functions of proteins have been discovered and new insights were gained^21–23^. In addition, based on the information of proteins with precise annotations and the relationships between proteins in the PPI networks, the functions of poorly functional characterized proteins can be inferred and the knowledge gaps can be filled^24,25^.

Several high-throughput experimental techniques such as yeast two-hybrid (Y2H), Tandem Affinity Purification and Mass Spectrometry (TAP-MS), protein chips, etc. have made significant contribution to the detection of PPIs for constructing genome-scale PPI networks^26–28^. However, these methods are intensive in labor and funds, let alone their inherent biases and limited coverage^29^. As far as we know, the number of PPIs detected by experimental methods are far from the estimated amount^30^. As a result, several types of *in silico* methods have been developed to meet the demand for determining missing PPIs in more species^31–33^. In this study, the protein interactome of *B. diazoefficiens* USDA110 was inferred by using “Interolog” and domain-based methods and reconstructed into a PPI network. The network was analyzed from different perspectives and compared with the PPI networks of other rhizobia. By integrating transcriptome and proteome data, PPI networks representing two typical physiological states - free-living (FL) cell and SNF bacteroid - were obtained and compared. Based on the SNF network, a core-sub-PPI-network related to symbiotic nitrogen fixation was determined and dissected, and novel functional modules and protein hubs were identified.

## 2. Materials and Methods

### 2.1. Generation of protein interactome

#### 2.1.1. Interolog method

According to the definition of “Interolog”^34^, for two proteins in *B. diazoefficiens* USDA110, if their homologous proteins in other species have an interaction, they are predicted to be interacting. For the “Interolog” method, we selected six prokaryotes as primary reference species including *Campylobacter jejuni*, *Escherichia coli*, *Helicobacter pylori*, *Mesorhizobium loti*, *Synechocystis sp. PCC6803* and *Treponema pallidum* (**Fig. 1**), all of which have relatively well-developed information of PPIs by experiments. PPIs of the six reference species were collected from literature^35–40^. All protein sequences of *B. diazoefficiens* USDA110 and of the six reference species were downloaded from NCBI RefSeq database. We predicted PPIs of *B. diazoefficiens* USDA110 by “Interolog” method in two ways. On one hand, protein sequences of the six reference species were prepared to do homologous alignment with *B. diazoefficiens* USDA110; on the other hand, experimental PPIs and their protein sequences from BioGRID^41^, DIP^42^, IntAct^43^ and HPRD_Release9_062910^44^ databases were added as supplement regardless of species. In order to improve the prediction accuracy, only the protein sequences from those databases which also appear in the Swiss-Prot database were used. The homologous proteins were acquired by using InParanoid^45^ (version 4.1) under default parameters.

**Figure 1.**
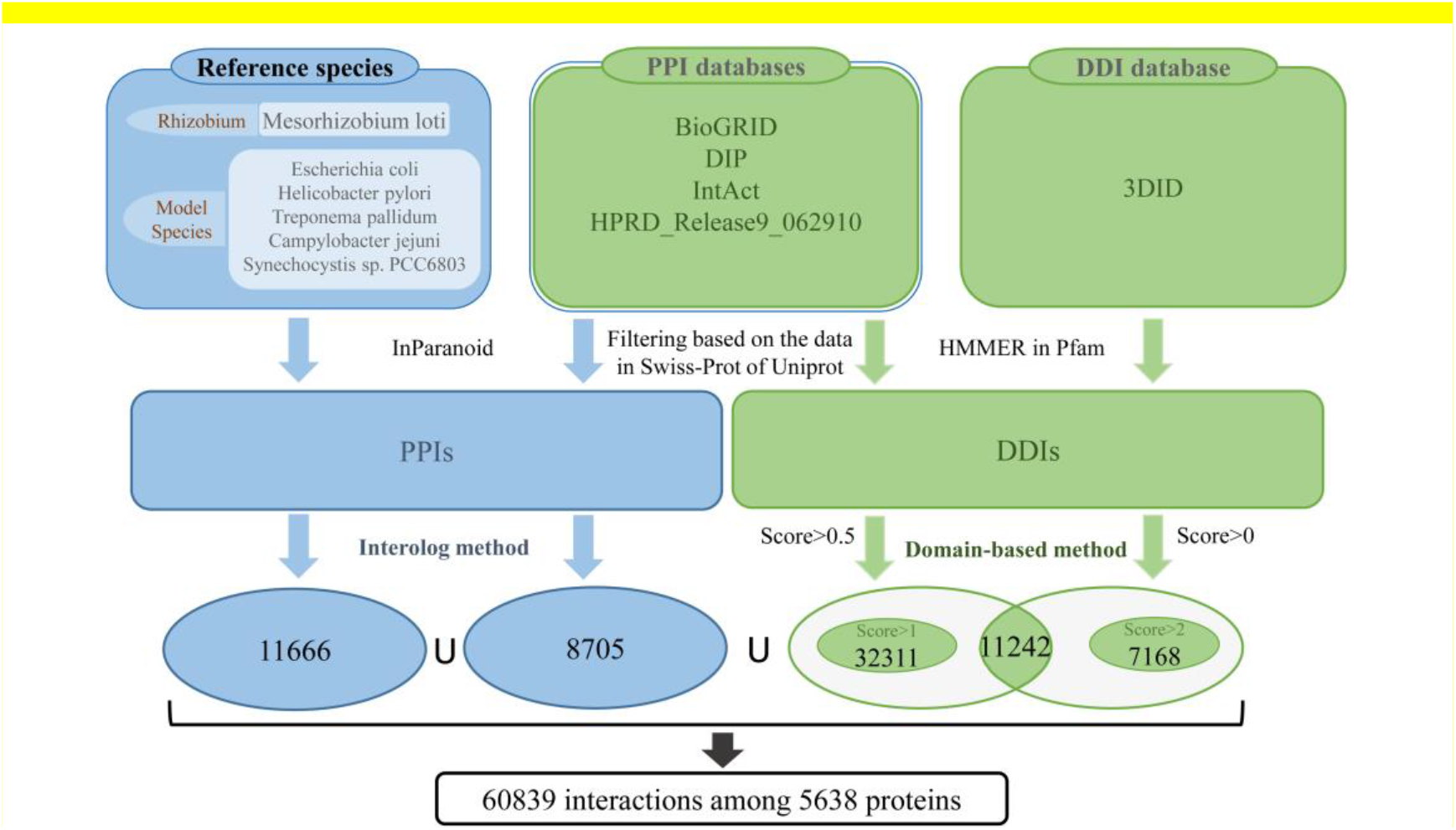
Flowchart for reconstructing the protein interactome of *B. diazoefficiens* USDA110.

#### 2.1.2. Domain-based method

In recent years, domain-based method has been widely used in PPI prediction, because the occurrence of PPI is usually caused by the interaction between one domain X of protein A and another domain Y of protein B and the interaction between proteins A and B can be inferred if domains X and Y have verified interaction. Here, domain-domain interactions (DDIs) were obtained at both the sequence and structure levels. The PPIs in databases of BioGRID, DIP, IntAct and HPRD_Release9_062910 (**Fig. 1**) were utilized to infer domain-domain interactions (DDIs). Besides, DDIs were also acquired from the three-dimensional interacting domains (3DID) database^46^. Protein domains in *B. diazoefficiens* USDA110 and in BioGRID, DIP, IntAct and HPRD_Release9_062910 databases were recognized and obtained based on the domain definitions of Pfam database and HMMER program (score>20, e-value<1e-5, coverage>0.9)^47,48^. A part of DDIs were inferred from experimental PPIs in databases based on domain sequences with (score>0.5) and another part of DDIs were obtained from 3DID database based on domain structure (score>0). As shown in **Fig. 1**, the intersection of these two parts as well as the domains of score>1 from PPI databases and the domains of score>2 from 3DID database were used in order to improve accuracy and data coverage.

### 2.2. Quality assessment of the reconstructed PPI network

First, the PPIs were verified by iLoop server (http://sbi.imim.es/iLoops.php) which utilizes local structure features to characterize protein interactions^49,50^. In our study, 1000 PPIs were randomly selected and submitted to the iLoop server for validation. Second, considering the interacting proteins are likely to appear in the same subcellular localization, the PPIs were verified by subcellular co-localization of the proteins. Sequences of the 5638 proteins in *B. diazoefficiens* USDA110 interactome were submitted to PSORT 3.0 (http://www.psort.org/psortb)^51^ and the location information of each protein was obtained. Third, the two proteins in a PPI tend to have similar function. Based on this point, the functional similarity of each interacting protein pair in the reconstructed PPI network and the randomized PPI network of the same topology was calculated according to a published algorithm G-SESAME^52^. The Gene Ontology (GO) function annotations of *B. diazoefficiens* USDA110 proteins were obtained from the GO database^53^ and Wilcoxon rank-sum test was performed to compare the functional similarity difference between the reconstructed PPI network and the random PPI network. Last, the correlation of transcriptional profiles between two genes from a PPI pair was also used to examine the reliability of the network. Here, the SNF sub-network of the global network were checked based on time-sequenced transcriptome data^11^. Pearson Correlation Coefficient (PCC) was calculated for the transcription profiles of every interacting pair in the reconstructed PPI network and the randomized PPI network of the same topology and then Wilcoxon rank-sum test was used to show the difference.

### 2.3. COG function enrichment analysis of PPIs

We assorted PPI numbers according to the Clusters of Orthologous Groups of proteins (COG) annotations as the functional network. Referred to 1000 random networks of the same topology generated from the functional network, the COG functional network was presented in the form of heat map. Z-scores were calculated to determine colors in the heat map by the following formula:

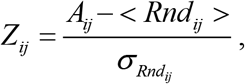

where *A*_*ij*_ represents the actual number of interacting protein pairs between COG category *i* and category *j*, < *Rnd*_*ij*_ > and 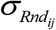 represent the mean and standard deviation of interacting protein pairs between category *i* and category *j* summarized from the 1000 random networks, respectively.

### 2.4. Validation of the PPI sub-networks in free-living and symbiotic nitrogen-fixing conditions

*B. diazoefficiens* USDA110 cells cultivated in aerobic peptone salts-yeast extract medium are considered as the FL condition. The processed microarray data of the FL *B. diazoefficiens* USDA110 were downloaded from NCBI-GEO database^54^ (GSM210269-GSM210283). In order to reduce false positive, a gene is deemed as expressed only in the case that more than 80% of the replicates are of *p*-value ≤ 0.06^11^. Then, the average of signals in selected replicates is regarded as the expression level^11^. The transcriptome and proteome data of bacteroids under the SNF state were obtained directly from the supplementary materials of a previous work^14^. The transcription level difference of an interacting protein pair *i* and *j* was normalized according to the following formula^55^:

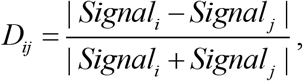

where *Signal*_*i*_ and *Signal*_*j*_ are the transcription levels of gene *i* and gene *j*, respectively.

## 3. Results and Discussion

### 3.1. Reconstruction of the genome-scale PPI network of *B. diazoefficiens* USDA110

The PPIs in *B. diazoefficiens* USDA110 were predicted by integration of an “Interolog” and a domain-based method (**Fig. 1**). The reconstructed PPI network was visualized by using Cytoscape 3.2.1^56^ (**Fig. S1A**), which contains 5638 proteins and 60839 PPIs. The Clusters of Orthologous Groups of proteins (COG) functional categories^57^ of the nodes in the PPI network were determined and their proportions are shown in **Fig. S1B**. Proteins related to ‘amino acid transport and metabolism (E)’ accounts for the largest proportion (over 12%), while the proteins associated with ‘cell cycle control, cell division, chromosome partitioning (D)’ only accounts for 0.5%.

### 3.2. Quality assessment of the PPI network

Reliability of the reconstructed PPI network was assessed from four different perspectives: local structural features, subcellular localization, functional similarities and transcriptional correlations of genes. For local structural features, 1000 PPIs were selected randomly and submitted to iLoop server by which the interactions were re-predicted. Due to the limitation of available structural templates (in the server’s dependent databases), 47.4% of the 1000 PPIs had at least one protein without corresponding structural features and thus cannot be assessed. For the remaining PPIs that can be assessed, 46.4% of the PPIs were confirmed, whereas only 1% was identified as non-interacting pairs (**Fig. 2A**), which proved that the reconstructed PPI network is reliable based on local structural features.

**Figure 2.**
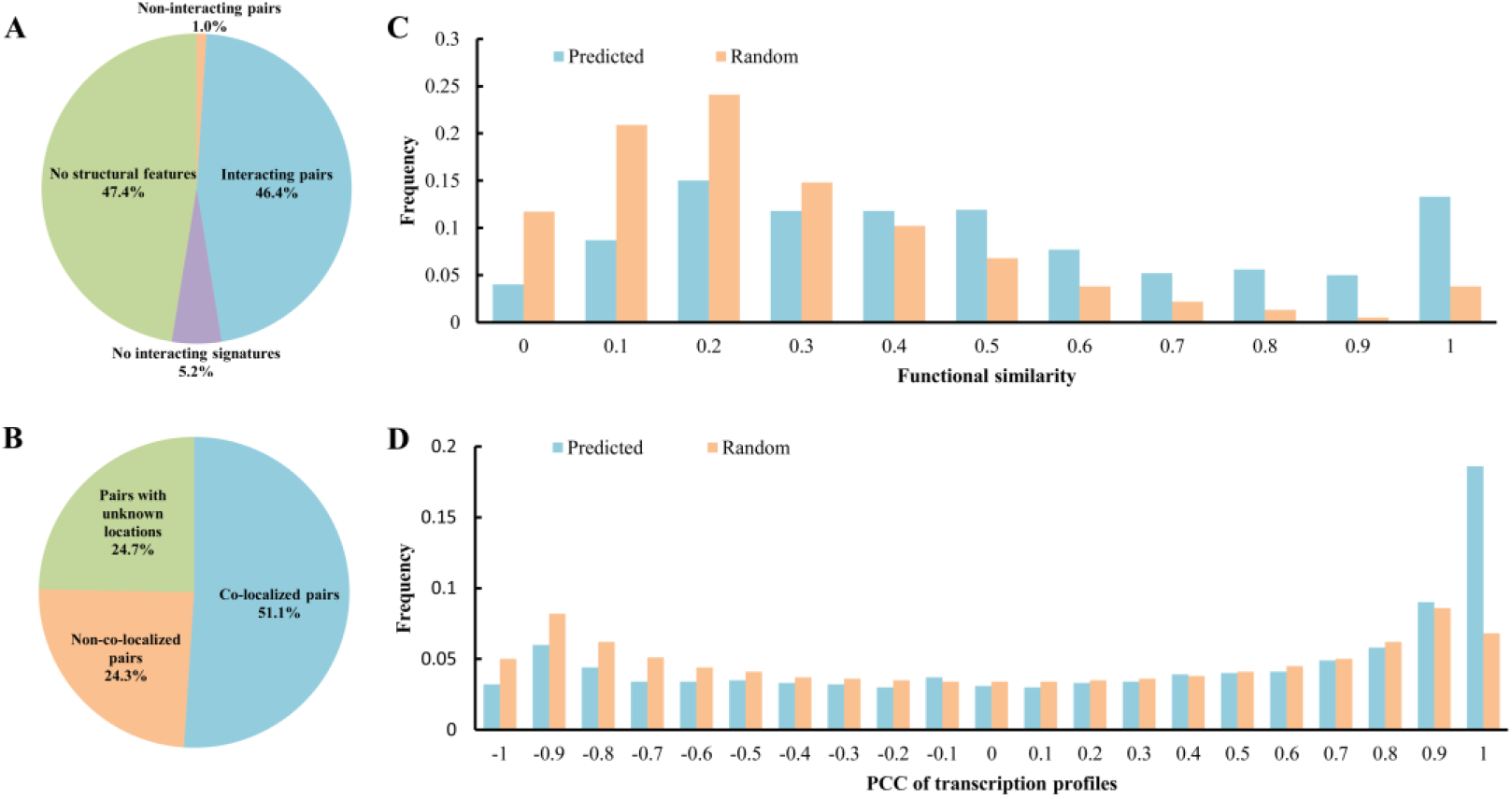
Validation of the *B. diazoefficiens* USDA110 protein interactome. (A) Validated by iLoop server. “Interacting pairs” represents the directly confirmed PPIs; “Non-interacting pairs” represents the directly non-confirmed PPIs; “No structural features” represents that no structural features (loop or domain) can be assigned to either one or both proteins; “No interaction signatures” represents that the pair of proteins have structural features but no interacting features. (B) Validated by subcellular co-localization. “Co-localized pairs” represents that two interacting proteins are co-localized; “Non-co-localized pairs” represents that two interacting proteins are not co-localized; “Pairs with unknown locations” represents that at least one protein in an interacting protein pair has no sub-cellular location information. (C) The difference of distribution of functional similarity between the reconstructed network and the randomized network of the same topology. Higher frequencies appear in the reconstructed network than the random network when functional similarity becomes larger. (D) The distribution difference of PCC of gene transcription profiles between the reconstructed network and the randomized network of the same topology. Higher frequencies appear in the reconstructed network than the random network when PCC of transcription profiles becomes larger.

The proteins which actually interact with each other are likely to appear in the same subcellular location. Here, we ascertained the subcellular location of the 5638 proteins in the reconstructed PPI network. According to the results, more than half of the PPIs (51.06%) were confirmed to be co-localized in the cell. Besides, over 25% PPI pairs have a component whose location is unknown (**Fig. 2B**), and for these PPI pairs some of them must be co-localized, therefore, the proportion of positive results is actually larger than 51.06%.

Previous studies have demonstrated that interacting proteins tend to have relatively higher functional similarities^37,58^, which can be used to confirm the reliability of the reconstructed PPI network. Based on the semantic similarities of gene ontology (GO) annotations^53^, the functional similarities of the proteins in PPI pairs of the reconstructed PPI network and corresponding random networks of the same topology were calculated and compared (**Fig. 2C**). It shows that the functional similarities in the reconstructed PPI network are significantly (*p*-value < 2.2e-16 from Wilcoxon rank-sum test) higher than those of the random networks, which proved that the reconstructed PPI network is reliable based on functional similarities of the proteins in PPI pairs.

It has been reported that interacting proteins have similar expression patterns^59^. Based on the temporal transcriptome profiles of the bacteroid of *B. diazoefficiens* USDA110^11^, the PCC of normalized transcription data of the proteins in PPI pairs of the reconstructed PPI network and corresponding random networks with the same topology were calculated and compared (**Fig. 2D**). The PCC values of the reconstructed PPI network are significantly (*p*-value < 2.2e-16 from Wilcoxon rank-sum test) higher than those of the random networks. Especially, when PCC is close to 1.0, the difference is more significant, which means that the reconstructed PPI network contains much more PPI pairs whose components keep fairly consistent transcriptional patterns than random networks. Hence, the reconstructed PPI network is reliable based on co-expression of genes.

### 3.3. Properties of the reconstructed PPI network

The topological parameters of the PPI network were calculated and analyzed with the “Network Analysis” plugin in Cytoscape^60^. Degree distribution of the PPI network of *B. diazoefficiens* USDA110 follows the power law (*y* = 2143.3*x*-1.348, *R*^2^=0.76, **Fig. 3A**), which shows the characteristic of scale-free network as many typical biological networks^61^. The number of sub-networks is 46 (**Fig. S1A**), and the largest sub-network contains 5556 proteins and 60730 interactions. The node degree distribution is shown in Fig. 3B. The distribution of cluster coefficient has two peaks (**Fig. 3C**), suggesting that the topological parameters of series of small sub-networks are very different from those of the largest one. The average shortest path length is around 3 and 4 (**Fig. 3D**), which is consistent with the results of previous studies^62^.

**Figure 3.**
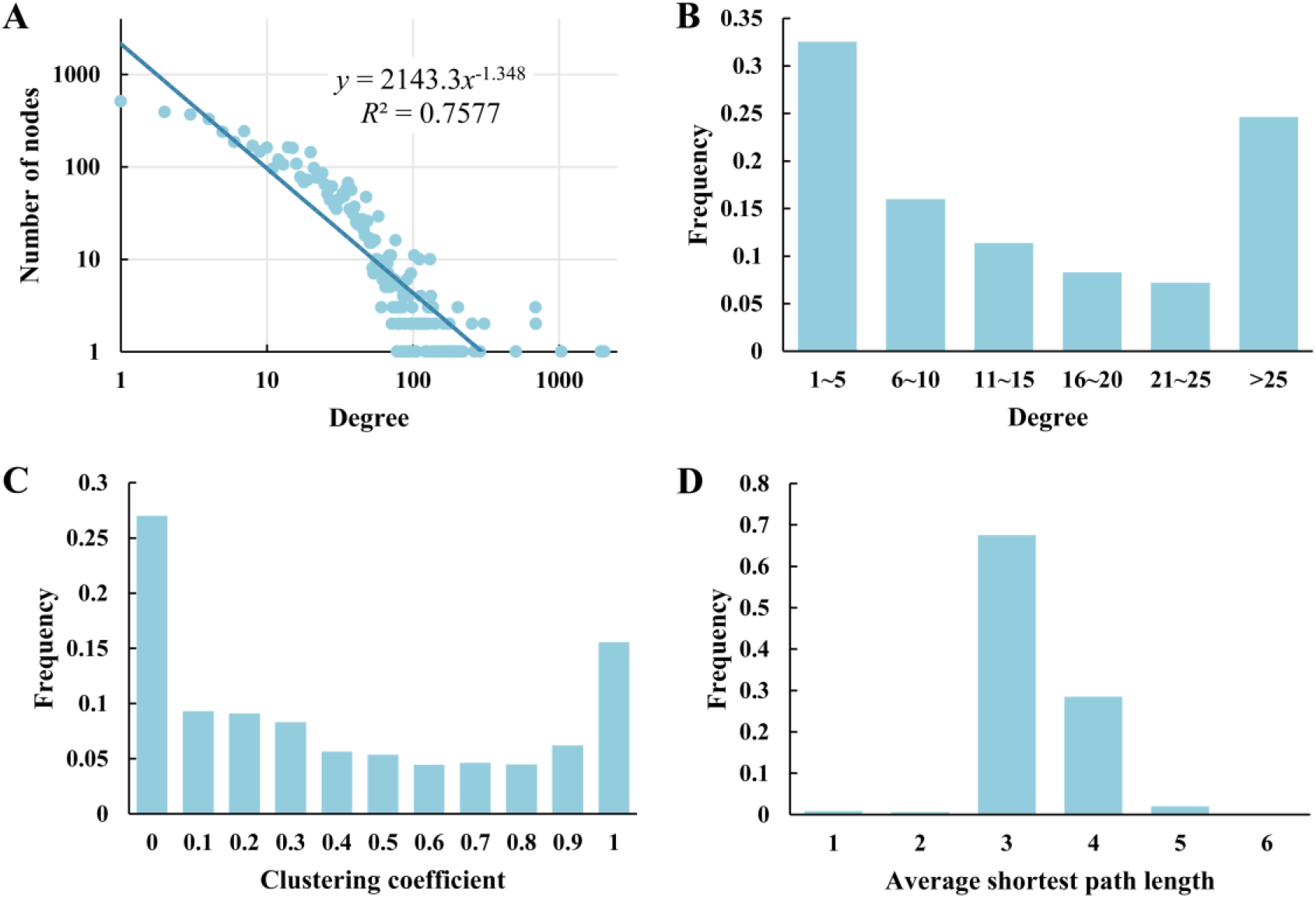
Distributions of topological properties of the *B. diazoefficiens* USDA110 PPI network.

In the power law fitting formula, the degree exponent *γ* of the reconstructed PPI network was calculated as 1.35 by the maximum likelihood estimate. Studies have shown that if the degree exponent is smaller than 2, the control of the whole network requires a small set of essential nodes which could be identified by minimum dominating set (MDS)^23,63^. By solving an integer-based linear programming problem^63^, a MDS of the *B. diazoefficiens* USDA110 PPI network was determined. In order to eliminate the bias caused by self-interactions, they were removed from the reconstructed PPI network, and the refined network (including 5611 proteins and 59125 PPIs) was used to identify the MDS. The determined MDS of the PPI network contains 427 nodes (less than 10% of the total number). Furthermore, COG enrichment analysis was performed to examine the functional distribution of these essential nodes. As shown in **Table S1**, the proteins in MDS are significantly enriched (Fisher’s exact test, *p* < 0.01) in ‘Energy production and conversion (C)’, ‘Lipid transport and metabolism (I)’, ‘Posttranslational modification, protein turnover, chaperones (O)’, ‘Intracellular trafficking, secretion, and vesicular transport (U)’ and ‘Defense mechanisms (V)’.

### 3.4. Comparison of PPIs among rhizobia

In the reconstructed PPI network of *B. diazoefficiens* USDA110, 5199 proteins with COG annotations (involving 55668 PPIs) were selected and analyzed. A z-score index which quantitatively reflects the degree of enrichment of PPIs in different combinations of COG categories was defined and calculated^40,64^. Same calculations were performed on the PPI networks of *M. loti* (2377 PPIs among 1408 proteins)^38^, *S. meliloti* (856 PPIs among 320 proteins)^65^ and *B. diazoefficiens* USDA110 under the SNF state (10473 PPIs among 1777 proteins). As shown in **Fig. 4**, proteins of the same functional categories are more likely to interact with each other in *B. diazoefficiens* USDA110 and *S. meliloti*, while this feature is not obvious in *M. loti*. Moreover, the PPIs related to the proteins of category O (posttranslational modification, protein turnover, chaperones) are the mainstay in the PPI network of *B. diazoefficiens* USDA110, because this protein category interacts with almost all the other protein categories. In contrast, the enrichment of PPIs related to category O in *M. loti* and *S. meliloti* is not obvious. This result may indicate that the PPIs in rhizobia are correlated with growth rate and the slow growth of *B. diazoefficiens* USDA110 requires more protein modification and repair that are performed by the proteins in category O. Another noticeable point is the interaction between protein categories J (translation, ribosomal structure and biogenesis) and U (intracellular trafficking, secretion, and vesicular transport); it is enriched in the *B. diazoefficiens* (SNF) and *S. meliloti* PPI networks which are both the sub-networks for symbiotic nitrogen fixation, while it is not so in *B. diazoefficiens* and *M. loti* which are both the global PPI networks. This result may suggest that protein biogenesis and cellular material transportation are correlated with each other through PPI in the SNF state in rhizobia, which enhances our understanding on the roles of material supply in the mechanism of symbiotic nitrogen fixation^66^.

**Figure 4.**
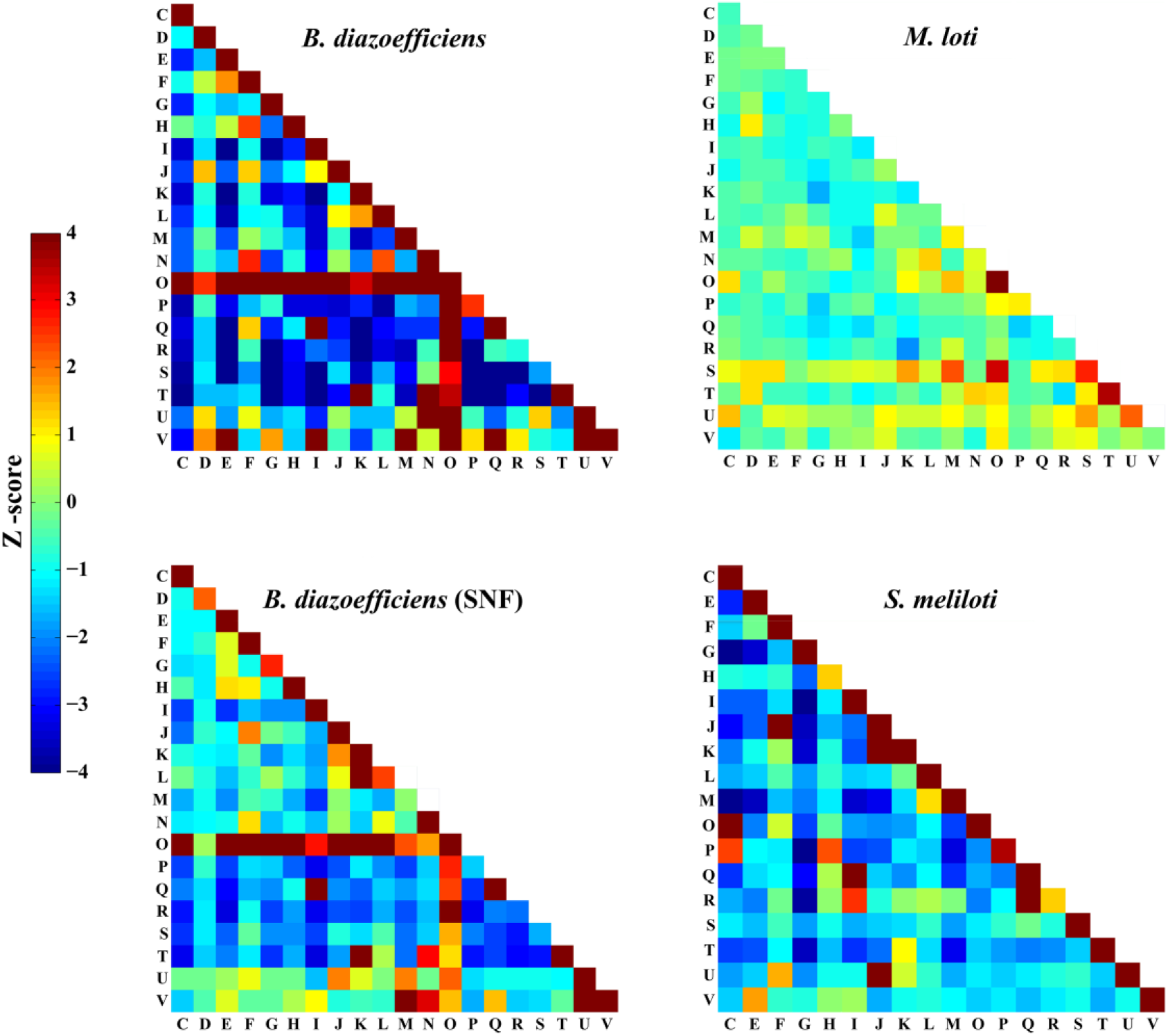
Enrichment in COG functional categories of PPIs for three organisms in different situations (the global PPI network: *B. diazoefficiens* USDA110, *M. loti*; the PPI network for SNF: *B. diazoefficiens* USDA110 (SNF), *S. meliloti*). The PPI numbers in COG categories were normalized by Z-score. Warmer color means higher enrichment.

The common functional modules such as signal pathways and protein complexes are relatively conservative among different species^67,68^. An algorithm called GASOLINE is available for the comparison of biological networks to find the conserved functional modules^69^. We used this algorithm to compare the PPI networks of several rhizobia for both the global network and the sub-network under SNF state. The global PPI networks of *B. diazoefficiens* USDA110 and *M. loti* have 26 pairs of conserved functional modules (Density threshold = 0.5, Sigma = 2) with particularly high-quality score (**Table S2**). For example, the largest module with the highest score (module 25 in **Table S2**, 6 proteins with ISC score 0.81) includes the hybrid sensors and regulators of two-component system for chemotaxis, and the protein interactions within this module and the correspondence relationships between the two compared species are shown in **Fig. 5A**. Since these species have special requirements for swimming in soil^70^, this module is considered to play a regulatory role in flagellum movement when rhizobia meet attractants in rhizosphere. As shown in **Fig. 5A**, 8 PPIs in this module match well between *B. diazoefficiens* USDA110 and *M. loti*. Considered the two additional PPIs (blr2194-bll1199, blr2192-bll1199) in *B. diazoefficiens* USDA110 and the correspondence relationship, it can be hypothesized that there is a chance that Q98HL7-Q98L88 and Q98PD0-Q98L88 in *M. loti* are also interacting protein pairs which might be missed in the experiments^38^.

**Figure 5.**
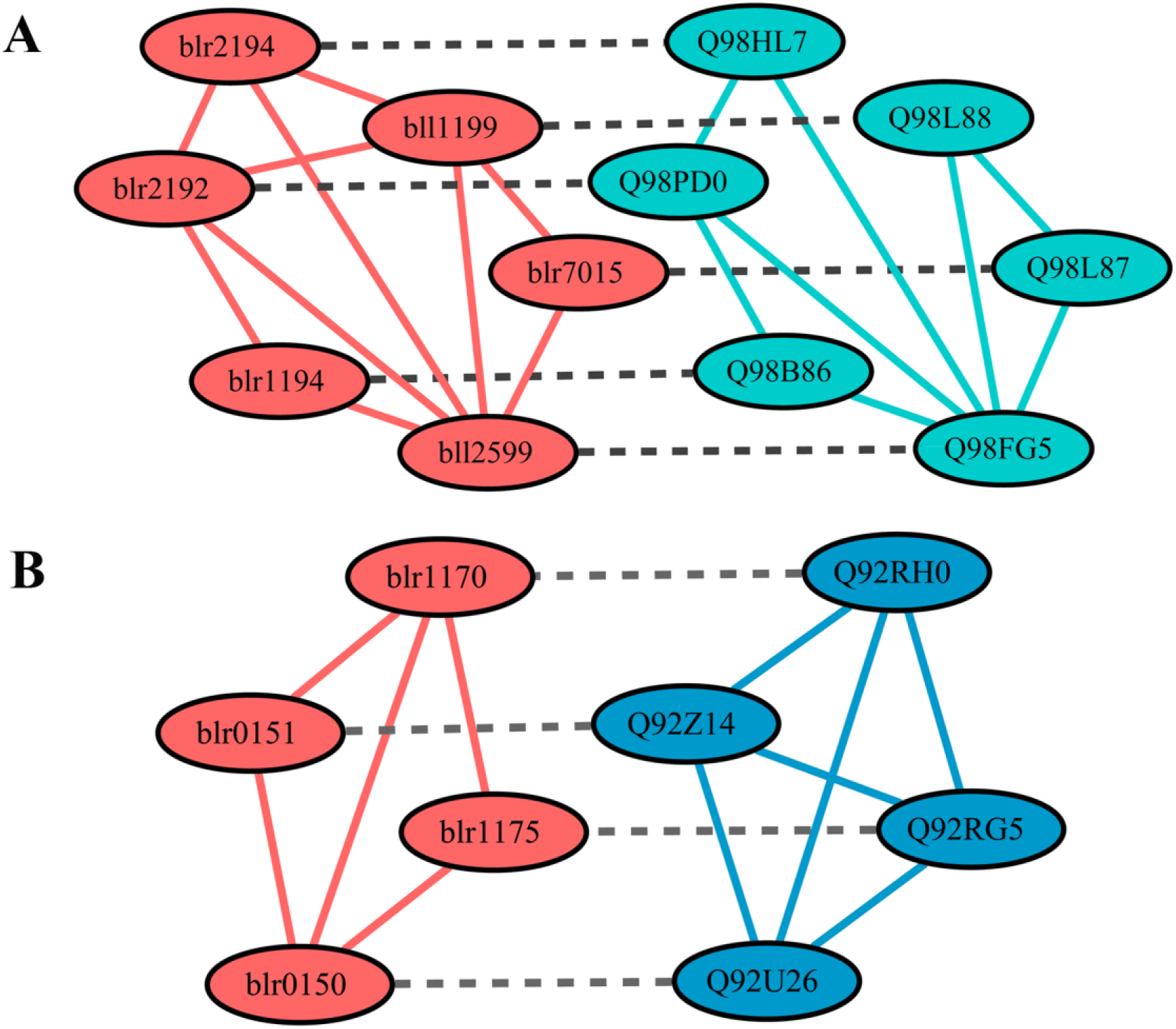
Examples of conserved PPI modules identified by GASOLINE. (A) The conserved PPI module related to chemotaxis was identified from the comparison between the global PPI network of *B. diazoefficiens* USDA110 (red) and M. loti (green). (B) The conserved PPI module related to cytochrome C oxidase was identified from the comparison between the SNF PPI network of *B. diazoefficiens* USDA110 (red) and *S. meliloti* (blue). Solid lines represent PPI within each module and dashed lines represent correspondence (homology) relationship between species.

For the SNF state, 10 pairs of conserved functional modules were obtained (Density threshold = 0.7,Sigma = 3) by comparing the PPI networks of *B. diazoefficiens* USDA110 and *S. meliloti* (**Table S3**). For example, the module with the highest score (module 1 in **Table S3**, 4 proteins with ISC score 0.88) includes ubiquinol-cytochrome enzyme (complex Ⅲ) subunit and cytochrome c oxidase (complex Ⅳ) subunit, and the protein interactions within this module and the correspondence relationships between the two species are shown in **Fig. 5B**. Since nitrogen fixation usually requires additional energy in terms of ATP and reductant, cytochrome c oxidase may produce the required ATPs to compensate the extra energy consumption^71^. Bhargava *et al* have proved that the activity of cytochrome c oxidase was enhanced in the process of nitrogen fixation^72^. The existence of the above PPI module (**Fig. 5B**) identified from *B. diazoefficiens* USDA110 and *S. meliloti* further validates the importance of cytochrome c oxidase in nitrogen fixation. Previous studies have shown that the process of electron transfer between complex Ⅲ and complex Ⅳ needs cytochrome c, but the specific process is still unknown^73^. In our study, the subunit components of complex Ⅲ (blr0151, blr0150; Q92Z14, Q92U26) and the subunit components of complex Ⅳ (blr1170, blr1175; Q92RH0, Q92RG5) all have interactions (**Fig. 5B**). Furthermore, in the 3DID database^46^, the domains (cox1, cox2, cox3) of subunits in complex Ⅲ and complex Ⅳ have been identified to be interacting. Based on these evidences, it can be inferred that complex Ⅲ and complex Ⅳ may have direct interaction under some states to improve the efficiency of electron transfer and release more required energy for nitrogen fixation.

In addition, with the reconstructed network of *B. diazoefficiens* USDA110, by network comparison with *M. loti* using GASOLINE and by consulting GO and KEGG databases, the functions of proteins belonging to COG category S (“function unknown”) were inferred (**Table S4**), which is a direct application of our reconstructed PPI network of *B. diazoefficiens* USDA110.

### 3.5. Analysis and comparison of the PPI networks under FL and SNF states

As the type strain of Bradyrhizobium rhizobia, *B. diazoefficiens* USDA110 has two typical physiological states: FL cell and SNF bacteroid. Therefore, it is meaningful to investigate the differences and similarities of the reconstructed PPI networks in these two states. Sub-networks representing the FL and SNF states were obtained by integrating transcriptome and proteome data^11,14^ into the global PPI network of *B. diazoefficiens* USDA110. The sub-network of FL state contains 3650 proteins and 31541 PPIs, while that of the SNF state contains 1777 proteins and 10473 PPIs. In order to compare the topological differences between these two networks, five local metrics were calculated for each node. Interestingly, although the proportion of common proteins in the FL and SNF networks exceeds 80% of the SNF network, all the topological metrics are significantly different. The results (**Table 1**) show that the median of nodes’ degrees is larger for the FL network due to its larger size and that all the other metrics suggesting the SNF network is more compact than the FL network, similar to the situation of the metabolic network of *B. diazoefficiens* USDA110^74^.

**Table 1.**
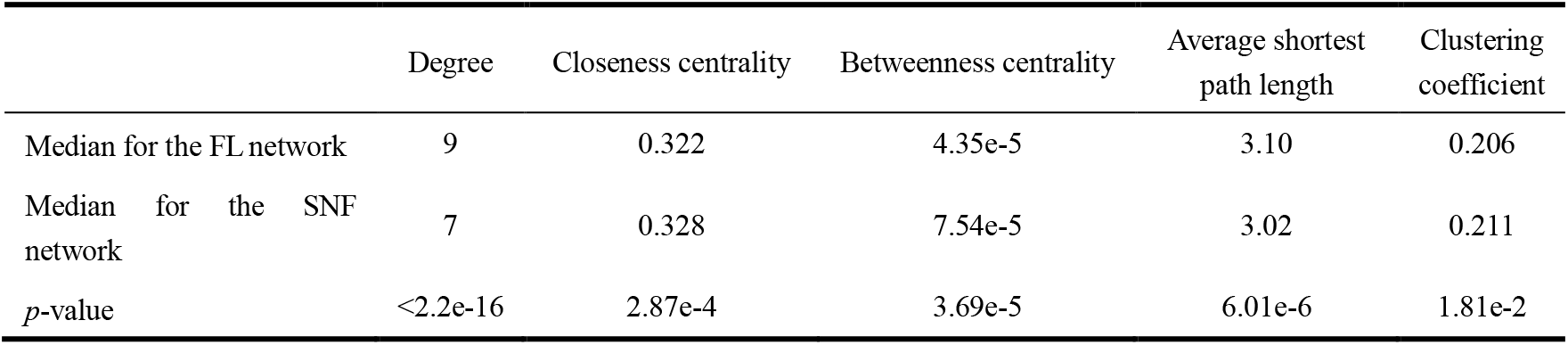
Comparison of local topological properties between the PPI networks of FL and SNF states (*p*-values were obtained by two-sided Wilcoxon rank sum test)

|  | Degree | Closeness centrality | Betweenness centrality | Average shortest path length | Clustering coefficient |
| --- | --- | --- | --- | --- | --- |
| Median for the FL network | 9 | 0.322 | 4.35e-5 | 3.10 | 0.206 |
| Median for the SNF network | 7 | 0.328 | 7.54e-5 | 3.02 | 0.211 |
| *p*-value | <2.2e-16 | 2.87e-4 | 3.69e-5 | 6.01e-6 | 1.81e-2 |

Previous studies have shown that the two components of interacting protein pairs in PPI network should have similar transcription levels^75^, which may reflect the trade-off between the pairing efficiency and synthesis cost. Therefore, the normalized transcription level difference between the two components of each protein pair in the PPI network was calculated and analyzed based on the transcriptome data of *B. diazoefficiens* USDA110^11,14,55^. Firstly, the distribution of normalized transcription difference of the protein pairs in FL and SNF networks (real PPIs, the “PPIs” group) was compared with that of all pairwise combinations of proteins in the corresponding network (all possible protein pairs, the “Control” group), respectively. As shown in **Fig. 6A**, the distribution of “PPIs” group is lower than that of “Control” group in both FL and SNF states (FL: *p*-value < 2.2e-16, median: 0.40 vs. 0.42, n: 31541 vs. 13322500; SNF: *p*-value = 3.42e-4, median: 0.37 vs. 0.38, n: 10473 vs. 3157729; compared by one-sided Wilcoxon rank sum test, *α* = 0.01), which means that the transcription levels of the interacting proteins are more consistent. Furthermore, similar comparisons were performed in each COG group in FL and SNF networks. The results showed that for the protein pairs belonging to the COG groups of C (energy production and conversion) and M (cell wall/membrane/envelope biogenesis), the normalized transcription differences of the FL state are significantly less than those of the SNF state, while for the COG groups of G (carbohydrate transport and metabolism), J (translation, ribosomal structure and biogenesis) and Q (secondary metabolites biosynthesis, transport and catabolism), the trend is just the opposite (**Fig. 6B**, **Table S5**).

**Figure 6.**
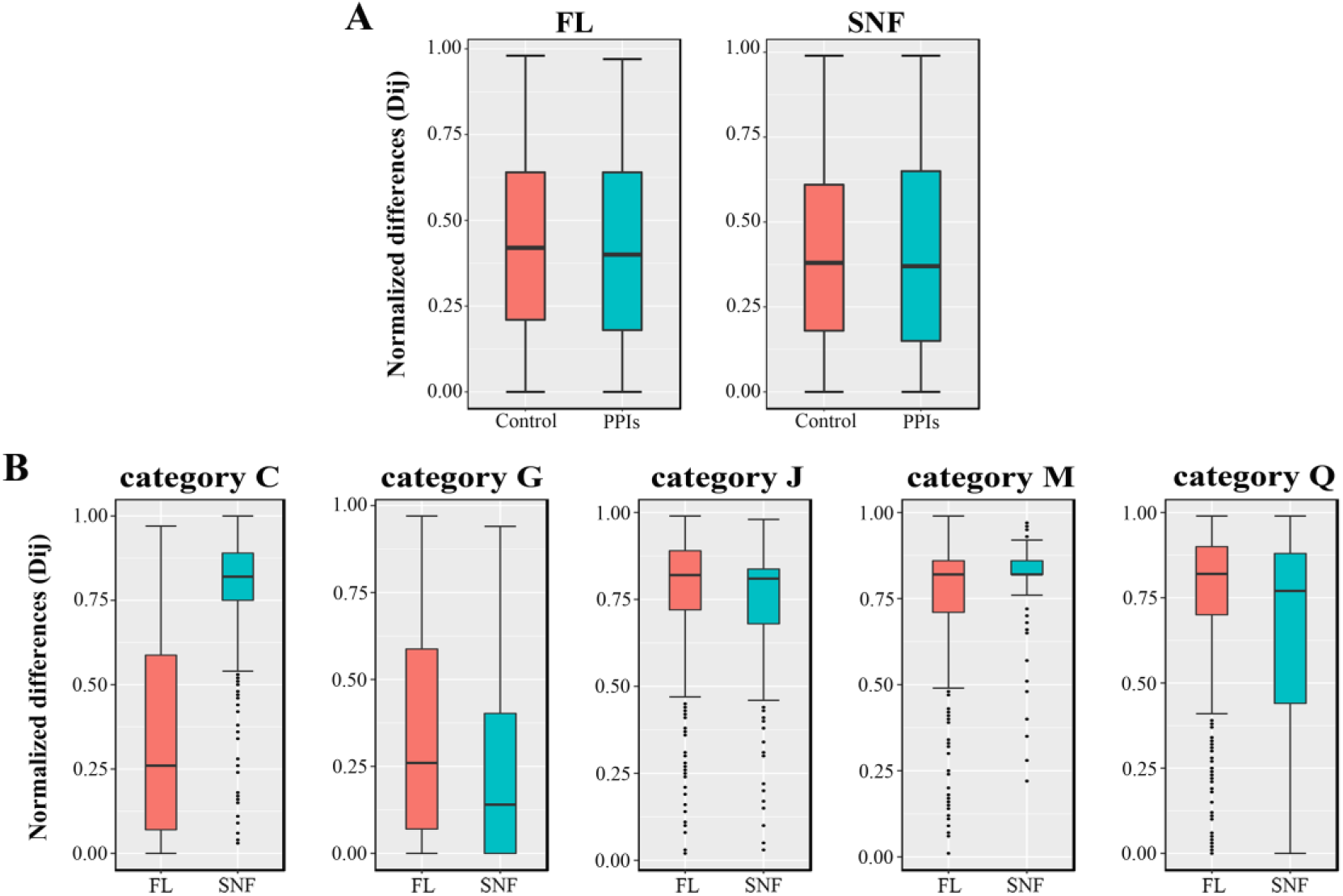
Comparison of normalized transcription level differences of interacting protein pairs. (A) Transcription differences in the Free-Living (FL) and Symbiotic Nitrogen Fixation (SNF) states. “Controls”, all pairwise combinations for the proteins in the reconstructed PPI network; “PPIs”, all the interacting protein pairs in the reconstructed PPI network. (B) Transcription differences in COG categories for the FL and SNF states. “category C”, the interacting protein pairs belonging to COG category C (energy production and conversion); and so on.

### 3.6. Analysis of the core-sub-PPI-network for symbiotic nitrogen fixation

Symbiotic nitrogen fixation is a complex physiological status involving many kinds of perfectly cooperated molecular machines. Functional genes related to symbiotic nitrogen fixation such as *nod*, *nif* and *fix* and their regulation have been a hotspot in the field of nitrogen-fixing bacteria, but their relationships with other physiological activities and relevant coordination are not very clear. Based on the reconstructed PPI network, functional modules of the core-sub-network related to SNF were identified and their cooperating relationships were analyzed.

Firstly, a dataset of 128 SNF-associated proteins in *B. diazoefficiens* USDA110 was determined by genome annotation and literature mining (**Table S6**). Based on these SNF-associated proteins, corresponding PPIs were filtered out and the SNF core-sub-network (SCSNW) was determined, which contains 195 proteins and 441 PPIs (**Supplementary file**). Using a network clustering tool based on the Markov chain clustering (MCL) algorithm in the clusterMaker plug-in for Cytoscape, called MCL Cluster^76,77^, a series of network modules of the SCSNW were obtained. As shown in **Fig. S2**, most of the SNF associated-proteins are hubs of modules, which validates that the modules identified by MCL clustering are functionally meaningful. Then nine key modules were selected out and their functions were deduced by gene annotations from different databases such as KEGG and UniProtKB as well as literature mining. As shown in **Table 2**, all of the modules are associated with different aspects of the phenotype of SNF bacteroid. Furthermore, the whole nitrogen fixing system also depends on the coordination of the modules which can be achieved through the PPIs between their components. In addition to these PPIs, functional modules can also be connected through other hubs, which can be named ‘Tie of Modules’ (TOM). Hubs with specific properties (**Fig. S3**) were selected out as TOMs. Through the 11 TOMs, more diversified coordination can be established between functional modules (**Fig. 7A**). Then the functions of PPIs binding TOMs and functional modules were deduced by genome annotation and literature mining (**Table S7**).

**Table 2.** Deduced functions of the network modules identified by MCL clustering

| Modules | No. of nodes | No. of PPIs | No. of SNF-associated proteins | Enriched COG categories | Deduced function |
| --- | --- | --- | --- | --- | --- |
| m1 | 24 | 49 | 2 | O, K, E, F | Synthesis of symbiotic nitrogen fixation associated protein |
| m2 | 18 | 40 | 4 | T | Oxygen concentration response and transcriptional regulation of downstream N2-fixing genes |
| m3 | 12 | 29 | 3 | E, C | Serine metabolism, transformation of amino - containing metabolites |
| m4 | 12 | 28 | 3 | T | Upstream regulation of nitrogenase and related transcription factors |
| m5 | 11 | 33 | 7 | E, C | N2-fixing reaction, NH<sub>4</sub><sup>+</sup> transport and related regulation |
| m6 | 11 | 32 | 4 | T | DNA damage repair, oxygen stress defense |
| m7 | 7 | 7 | 1 | E | Transformation of amino - containing metabolites |
| m8 | 7 | 11 | 4 | C | Energy production, electron transfer |
| m9 | 6 | 8 | 2 | C | Reduction of oxidative nitrogenase |

**Figure 7.**
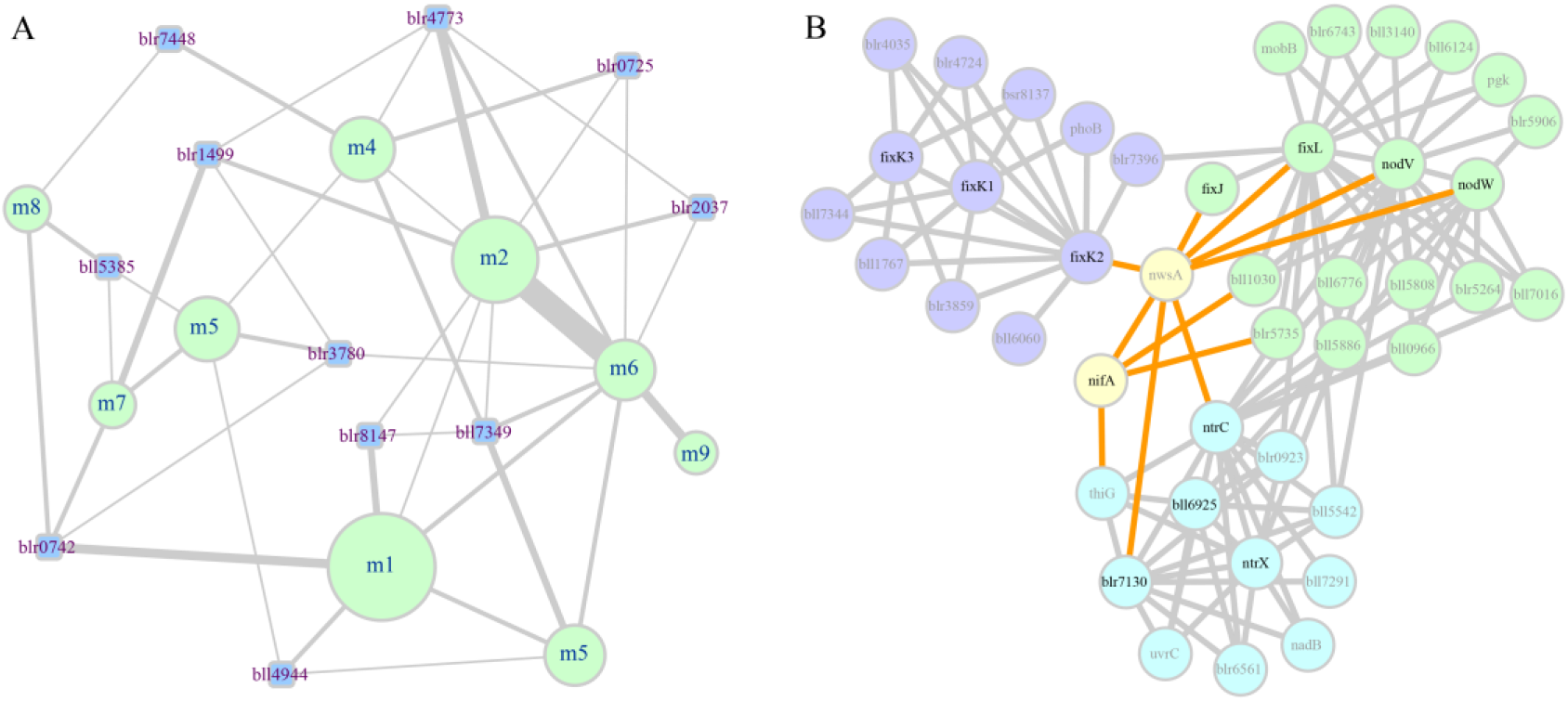
Sub-networks related to key modules (MCL clusters) and TOMs in the SNF core-sub-network. (A) Sub-network related to 9 key modules (light-green nodes) and 11 TOMs (light-blue nodes). The diameter of a key module node corresponds to the number of proteins in this module. The width of an edge between two nodes corresponds to the number of PPIs. (B) Sub-network related to NwsA (including its interacting modules and TOMs). Light-green, blue and purple nodes represent the m2, m4 and m6 modules, respectively. Nodes with black labels represent the SNF-associated proteins.

As shown in **Fig. 7B**, we found protein NwsA, a two-component hybrid sensor and regulator coded by gene blr4773, connects three functional modules (m2, m4, m6) and interacts with several key hubs of regulatory cascades in SNF bacteroid. As to these regulators, NodV and NodW regulate the expression of other nod genes (such as nodYABC) that initiate nodulation^78,79^; FixLJ-FixK2-FixK1 cascade activates genes essential for microoxic respiration in symbiosis (such as fixNOPQ and fixGHIS) and further regulatory genes for nitrogen fixation (rpoN1, nnrR, and fixK1)^80–82^; NifA controls expression of nitrogen fixation genes (such as nifHDK)^81,83^; NtrC partly controls the regulation of genes related to nitrogen metabolism (such as ginII)^84,85^. Studies have reported that cross talk between the FixLJ-FixK2-FixK1 cascade and RegSR-NifA cascade might allow switching between different expression patterns of genes essential for N2 fixation^81,86^. Therefore, NwsA might play a critical role in the cross talk and probably perform an important coordination function between nodulation, nitrogen fixation and nitrogen assimilation in symbiotic nitrogen fixation system, which waits for experimental validation. Just like NwsA, other TOMs might also be important for symbiotic nitrogen fixation through interactions with related proteins, although further information and knowledge need to be accumulated.

## 4. Conclusions

In this work, the protein interactome (60839 PPIs among 5638 proteins) of *B. diazoefficiens* USDA110 was computationally reconstructed by combining Interolog and domain-based methods and its reliability was validated from four perspectives. The reconstructed interactome of *B. diazoefficiens* USDA110 was compared with those of other rhizobia (*M. loti* and *S. meliloti*) and it was inferred that the slow growth of *B. diazoefficiens* USDA110 may require more protein modification and repair via protein-protein interaction. By network comparison, 36 conserved functional modules were identified and the functions of related proteins were annotated. By integrating the reconstructed interactome with transcriptome and proteome data, the sub-networks representing FL and SNF states of *B. diazoefficiens* USDA110 were derived. Based on the SNF sub-network and the SNF-associated proteins, 9 novel functional modules and 11 protein hubs (named TOMs which connect functional modules) were identified and analyzed to further our understanding on the molecular mechanism of symbiotic nitrogen fixation.

## Abbreviations

PPI: protein-protein interaction
SNF: symbiotic nitrogen-fixing
FL: free-living
DDI: domain-domain interaction
GO: gene ontology
PCC: Pearson correlation coefficient
COG: clusters of orthologous groups of proteins
GEO: gene expression omnibus
MDS: minimum dominating set
3DID: three-dimensional interacting domains
MCL: Markov chain clustering
TOM: Tie of Modules
SCSNW: the SNF core-sub-network

## Acknowledgements

This work was supported by the National Natural Science Foundation of China (Grant 31971184), Project 2662016PY094 supported by the Fundamental Research Funds for the Central Universities, and the National Basic Research Program of China (973 project, Grant 2013CB127103). The funders had no role in study design, data collection and interpretation, or the decision to submit the work for publication. We thank Bao-Hai Hao and Youguo Li for valuable discussion.

## Author Contributions

Conceived and designed the experiments: B.G.M. Performed the experiments: J.X.M., G.L., Y.Y. Analyzed the data: J.X.M., Y.Y. B.G.M. Wrote the paper: J.X.M., Y.Y. and B.G.M.

## Additional Information

**Supplementary information** accompanies this paper at http://

