## Supplementary information for "Computationally reconstructed interactome of *Bradyrhizobium diazoefficiens* USDA110 reveals novel functional modules and protein hubs for symbiotic nitrogen fixation"

Supplementary Contents:

Supplementary Figures S1-S3.

Supplementary Table S1-S7.

Supplementary file.xlsx

### Supplementary information

#### Supplementary figures

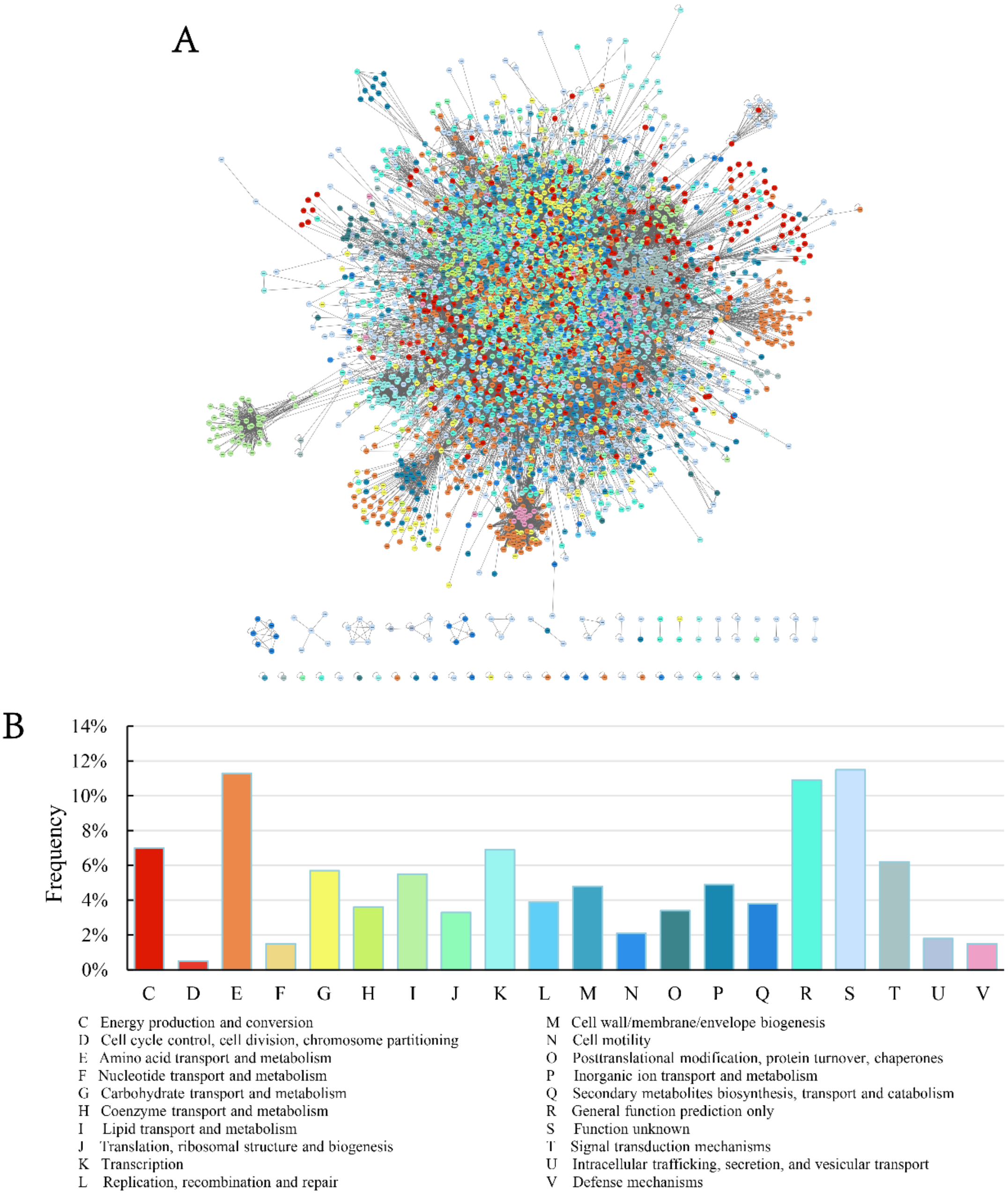


**Fig. S1** (A) The reconstructed protein-protein interaction (PPI) network of *B. diazoefficiens* USDA110. (B) The proportions of COG functional categories in the PPI network. The nodes in the PPI network were colored in accordance with COG functional categories.

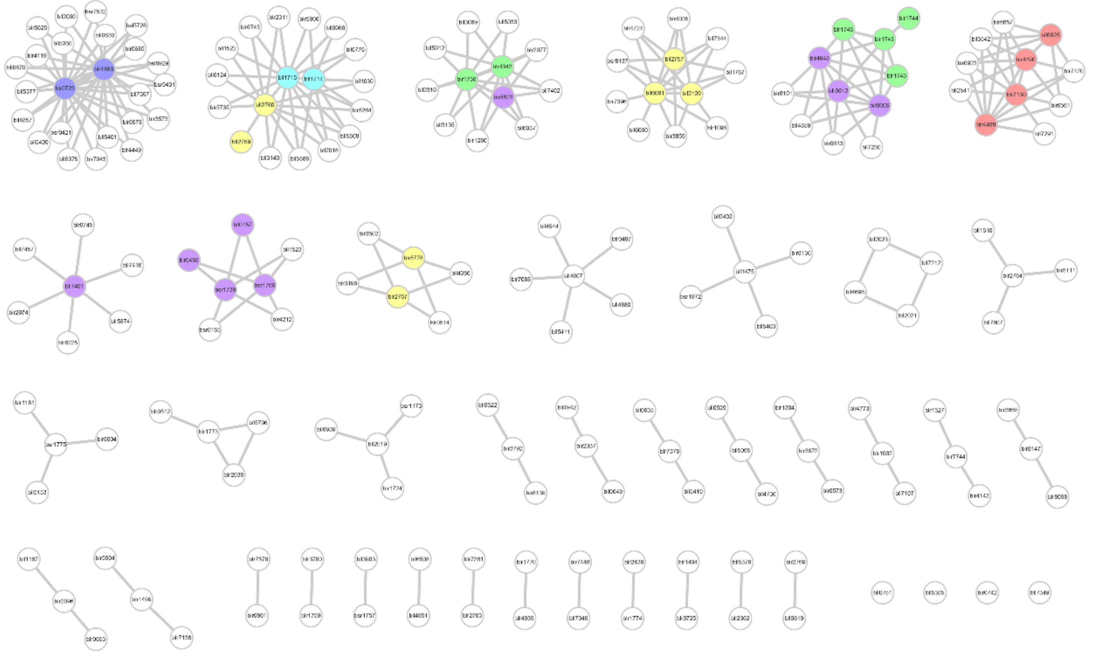

**Fig. S2** Functional modules of SNF core-sub-network in the PPI network of *B. diazoefficiens* USDA110 identified by MCL clustering. The modules were arranged in descending order by module size and the first 9 modules (m1-m9) were colored. The node colors represent different kinds of SNF-associated proteins: purple, sigma54 factor; cyan, nod proteins; yellow, fix proteins; green, nif proteins; red, two component regulators; violet, other SNF-associated proteins.

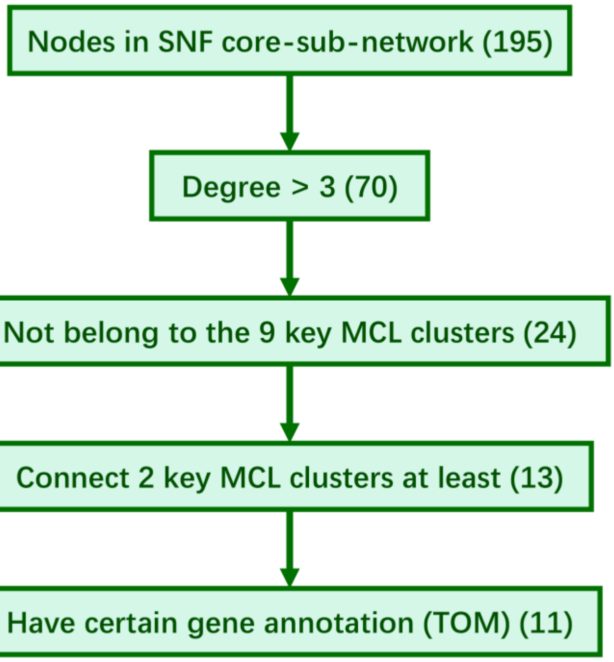


**Fig. S3** Schematic representation of the procedure used to select hubs with specific properties named Tie of Modules (TOM) in the SNF core-sub-network of *B. diazoefficiens* USDA110 PPI network.

### Supplementary tables

**Table S1. Significantly enriched COG categories in the MDS of the reconstructed PPI network (fisher's exact test, *p*-value<0.01)**

| COG category | Number of proteins in PPI network | Number of proteins in MDS | *p*-value |
|---|---|---|---|
| V Defense mechanisms | 96 | 6 | 6.53e-6 |
| I Lipid transport and metabolism | 349 | 8 | 1.49 e-4 |
| O Posttranslational modification, protein turnover, chaperones | 210 | 25 | 2.01e-3 |
| U Intracellular trafficking, secretion, and vesicular transport | 61 | 11 | 7.78 e-3 |
| C Energy production and conversion | 414 | 37 | 9.72 e-3 |

**Table S2. Conserved functional modules of the global PPI networks of *B. diazoefficiens* USDA110 and *M. loti* obtained by the GASOLINE algorithm**

| Module | *B. diazoefficiens* USDA110 | *M. loti* | ISC score | Module size |
|---|---|---|---|---|
| 1 | blr3189,bll7021,blr7895,blr3104 | Q98IV9,Q98BL5,Q98DQ2,Q98M07 | 1.00 | 4 |
| 2 | bll1799,bll8025,bll1317,bll3903 | Q989N3,Q98ID9,Q98AS5,Q98B05 | 1.00 | 4 |
| 3 | blr4439,bll2403,bll5459,bll7190 | Q98K28,Q98H78,Q98K67,Q987J9 | 1.00 | 4 |
| 4 | bll4303,bll0933,blr1822,bll3903 | Q98GT2,Q983Q0,Q989N6,Q98B05 | 1.00 | 4 |
| 5 | blr7523,bll7180,bll2795,blr4388,bll2590 | Q987U8,Q98DA7,Q98P62,Q98JF1,Q988U1 | 0.96 | 5 |
| 6 | blr3074,bll6508,bll3903,blr4773,blr6751 | Q985J1,Q98KI9,Q98B05,Q98LK8,Q98IT8 | 0.96 | 5 |
| 7 | bll6164,bll6258,bll8025,bll3903,blr3064 | Q98CI8,Q983P9,Q98ID9,Q98B05,Q98GJ6 | 0.96 | 5 |
| 8 | bll5925,blr0616,blr8186,bll3903,bsl1473 | Q985Q9,Q98EU9,Q98KE1,Q98B05,Q98PA0 | 0.96 | 5 |
| 9 | bll3903,bll5925,blr2027,blr6216 | Q98B05,Q985Q9,P17862,Q98LU4 | 0.94 | 4 |
| 10 | bll7306,bll2759,bll2760,bsl5082 | Q98DH9,Q988T4,Q988T3,Q988T5 | 0.94 | 4 |
| 11 | blr7730,blr2168,blr4157,bll3551 | Q983H9,Q98B19,Q98ME0,Q986X5 | 0.94 | 4 |
| 12 | bll3246,bll6292,bsr6998,bll6711 | Q98FP1,Q98MG5,Q98HL2,Q987J3 | 0.94 | 4 |
| 13 | blr2027,blr6216,bll5925,bll3903 | P17862,Q98LU4,Q985Q9,Q98B05 | 0.94 | 4 |
| 14 | blr0425,bll6773,bll3903,bll1317 | Q98EZ3,Q98P39,Q98B05,Q98AS5 | 0.94 | 4 |
| 15 | bll0465,blr0616,bll0109,blr8186 | Q98EA1,Q98EU9,Q98L85,Q98KE1 | 0.94 | 4 |
| 16 | bll8024,bll0109,bll5913,bll3903,blr6172 | Q98B49,Q98L85,Q985K2,Q98B05,Q98GX4 | 0.92 | 5 |
| 17 | blr5104,blr5132,bll1799,blr7359,bll4958 | Q98M83,Q98MA8,Q989N3,Q98E49,Q989W9 | 0.92 | 5 |
| 18 | bll0378,blr5523,bll5913,bll3903,blr6753 | Q986V5,Q98LQ7,Q985K2,Q98B05,Q98IU1 | 0.92 | 5 |
| 19 | bll2590,bll2795,bll0109,blr4388,bsl0578 | Q988U1,Q98P62,Q98L85,Q98JF1,Q98ET2 | 0.92 | 5 |
| 20 | bll0457,bll6508,bll4898,bll0109,blr6172 | Q98EC2,Q98KI9,Q986M8,Q98L85,Q98GX4 | 0.92 | 5 |
| 21 | blr1228,blr4388,blr0711,bll0109,blr3064 | Q98CW4,Q98JF1,Q98CP4,Q98L85,Q98GJ6 | 0.92 | 5 |
| 22 | bll1475,bll0109,blr3873,bll5663,blr1095 | Q985Q4,Q98L85,Q98K79,Q98NL5,Q98BC0 | 0.88 | 5 |
| 23 | bll0109,blr6774,bll4898,bll5663,bll1475 | Q98L85,Q98DP7,Q986M8,Q98NL5,Q985Q4 | 0.84 | 5 |

| | | | | |
|---|---|---|---|---|
| 24 | blr0329,bll0659,bll3140,blr3467 | Q98L88,Q98FG5,Q985F5,Q98L87 | 0.81 | 4 |
| 25 | blr1194,bll2599,bll1199,blr7015,blr2192,blr2194 | Q98B86,Q98FG5,Q98L88,Q98L87,Q98PD0,Q98HL7 | 0.81 | 6 |
| 26 | bll4573,blr2192,bll0659,blr6651,blr2194,bll3706 | Q98HI5,Q98PD0,Q98FG5,Q98E43,Q98HL7,Q98B86 | 0.78 | 6 |

**Table S3. Conserved functional modules of the PPI networks of *B. diazoefficiens* USDA110 and *S. meliloti* under SNF state obtained by the GASOLINE algorithm**

| Module | *B. diazoefficiens* USDA110 | *S. meliloti* | ISC score | Module size |
|---|---|---|---|---|
| 1 | blr1170,blr0150,blr1175,blr0151 | Q92RH0,Q92U26,Q92RG5,Q92Z14 | 0.88 | 4 |
| 2 | bll6899,blr3345,blr5675,bll4543 | Q92RF4,Q92MM9,Q92MN0,Q92MM8 | 0.75 | 4 |
| 3 | blr5675,bll6899,bll4542,bll4543 | Q926C5,Q92MN0,Q92MX4,Q92MX3 | 0.75 | 4 |
| 4 | blr3725,bll3808,blr2146,blr0770 | P50205,P06230,P06234,Q92SV9 | 0.69 | 4 |
| 5 | blr2146,bll3808,blr0770,blr3725 | P06234,Q92P54,P06230,P50205 | 0.69 | 4 |
| 6 | blr4490,blr4488,blr7130,bll7291 | Q92Q87,P13632,P10577,P03028 | 0.69 | 4 |
| 7 | blr4488,blr5735,blr5264,bll5808,bll0966,blr3071 | P10577,P03028,Q92Q87,Q52977,P13632,Q92TC2 | 0.67 | 6 |
| 8 | bll3809,blr3725,bll3808,blr0770 | P56902,P06234,P06230,Q92SV9 | 0.63 | 4 |
| 9 | blr6078,blr5675,bll7146,bll6899 | Q92MN0,Q92ZZ6,Q92MM9,Q926C5 | 0.63 | 4 |
| 10 | blr3353,bll7146,bll6899,blr5675 | Q92ZZ6,Q92MM9,Q92MN0,Q926C5 | 0.63 | 4 |

**Table S4. Functional annotation of the proteins in COG category S ("function unknown") by using the conserved PPI modules in *B. diazoefficiens* USDA110 and *M. loti* (network comparison was performed by using the GASOLINE algorithm; functions are inferred based on interacting partners and GO and KEGG annotations; The highlighted genes have no annotations in GO and KEGG databases and purely predicted by the PPI networks)**

| *B. diazoefficiens* USDA110 | COG | Annotation of gene function | *M. loti* | COG | Annotation of gene function |
|---|---|---|---|---|---|
| bll8024 | S | ATP binding | Q98B49 | S | ATP binding |
| blr6172 | S | cellular response to DNA damage stimulus | Q98GX4 | S | cellular response to DNA damage stimulus |
| blr6753 | S | transcriptional regulator related to branched-chain amino acid biosynthesis | Q98IU1 | S | transcriptional regulator related to branched-chain amino acid biosynthesis |
| blr6774 | S | double-stranded DNA binding | Q98DP7 | S | double-strand break repair via nonhomologous end joining |
| blr3189 | S | integral component of membrane | Q98IV9 | S | integral component of membrane |
| blr7895 | S | metal ion binding；<br>oxidoreductase activity | Q98DQ2 | S | metal ion binding；<br>oxidoreductase activity |

| | | | | | |
|---|---|---|---|---|---|
| blr6216 | S | formaldehyde catabolic process | Q98LU4 | S | formaldehyde catabolic process |
| bll0465 | S | peptidase related to abruption of rhizobia | Q98EA1 | S | peptidase related to abruption of rhizobia |
| blr8186 | S | transferase activity | Q98KE1 | S | transferase activity |
| blr7523 | S | 16S ribosomal RNA methyltransferase RsmE | Q987U8 | S | 16S ribosomal RNA methyltransferase RsmE |
| blr3074 | - | the process of coordination between nitrogen fixation, nodulation and nitrogen assimilation | Q985J1 | - | the process of coordination between nitrogen fixation, nodulation and nitrogen assimilation |
| blr6751 | S | response to stress | Q98IT8 | S | response to stress |
| blr8186 | S | transferase activity | Q98KE1 | S | transferase activity |
| bsl1473 | S | transcriptional regulator related to abruption of rhizobia | Q98PA0 | S | transcriptional regulator related to abruption of rhizobia |
| bsl0578 | S | peptidase related to amino acid metabolism | Q98ET2 | S | peptidase related to amino acid metabolism |
| blr6172 | S | cellular response to DNA damage stimulus | Q98GX4 | S | cellular response to DNA damage stimulus |

**Table S5. Comparison of normalized transcription level differences of the interacting proteins belonging to the same COG category under FL and SNF states (the groups with significant differences are shown, one-sided Wilcoxon rank sum test, $\alpha$ = 0.05)**

| COG category | No. of PPIs under FL state | No. of PPIs under SNF state | $D_{ij}$ Median under FL state | $D_{ij}$ Median under SNF state | *p*-value |
|---|---|---|---|---|---|
| Energy production and conversion (C) | 638 | 349 | 0.26 | 0.82 | <2.20e-16 |
| Carbohydrate transport and metabolism (G) | 490 | 116 | 0.35 | 0.14 | 6.48e-8 |
| Translation, ribosomal structure and biogenesis (J) | 421 | 166 | 0.82 | 0.81 | 2.85e-3 |
| Cell wall/membrane/ envelope biogenesis (M) | 329 | 89 | 0.82 | 0.82 | 0.0152 |
| Secondary metabolites biosynthesis, transport and catabolism (Q) | 519 | 239 | 0.82 | 0.77 | 1.29e-6 |

**Table S6. The 128 SNF-associated proteins in *B. diazoefficiens* USDA110 determined by genome annotation and literature mining**

| SNF-associated proteins in *B. diazoefficiens* USDA110 | | | | | | | | |
|---|---|---|---|---|---|---|---|---|
| bll0157 | bll0416 | blr0488 | blr0517 | blr0606 | blr0612 | blr0653 | blr0680 | blr0723 |
| blr0724 | blr0725 | blr0742 | bll0800 | blr0906 | blr1062 | blr1063 | bll1167 | blr1383 |
| bll1421 | bll1475 | blr1499 | bll1630 | bll1631 | blr1632 | bll1714 | bll1715 | bsr1739 |
| blr1743 | blr1744 | blr1745 | blr1746 | blr1755 | blr1756 | bsr1757 | blr1759 | bsr1760 |

| | | | | | | | | |
|---|---|---|---|---|---|---|---|---|
| **blr1769** | **blr1773** | **blr1774** | **bsr1775** | **blr1813** | **blr1815** | **blr1883** | **blr1902** | **bll2019** |
| **bll2021** | **bll2023** | **blr2026** | **blr2027** | **blr2028** | **blr2029** | **blr2030** | **blr2031** | **blr2033** |
| **blr2034** | **blr2036** | **blr2037** | **blr2038** | **blr2062** | **bll2109** | **bll2362** | **blr2369** | **bll2623** |
| **blr2694** | **bll2757** | **bll2759** | **bll2760** | **blr2763** | **blr2764** | **blr2766** | **blr2767** | **blr2769** |
| **blr2807** | **blr3125** | **blr3126** | **blr3127** | **blr3128** | **bll3460** | **bll3466** | **blr3792** | **blr3959** |
| **blr3972** | **blr4338** | **blr4342** | **blr4486** | **blr4487** | **blr4488** | **blr4489** | **blr4490** | **bll4607** |
| **bll4612** | **blr4614** | **blr4948** | **bll4951** | **bll4952** | **bll5061** | **bll5065** | **bll5464** | **bll5465** |
| **bll5476** | **blr5523** | **blr5778** | **bll6061** | **bll6925** | **blr7003** | **blr7130** | **bll7191** | **bll7193** |
| **blr7378** | **bll7401** | **blr7496** | **blr7573** | **bll7574** | **blr7577** | **blr7578** | **bll7696** | **blr7744** |
| **bll7861** | **blr7884** | **blr7900** | **bll7944** | **bll7945** | **bll7946** | **bll7947** | **blr8147** | **blr2768** |
| **blr1770** | **blr1771** | | | | | | | |

**Table S7. Deduced functions of TOMs (Tie of Modules)**

| Synonym | Gene | COG | SNF associated proteins or not | Degree | Connected modules and TOMs | Annotation of gene function | Deduced function of related PPIs | Refs |
|---|---|---|---|---|---|---|---|---|
| bll7349 | sigA | K | N | 11 | m2 m4 m5 m6 blr8147 | RNA polymerase sigma factor RpoD | Regulating binding strength of sigma factor RpoD to its binding sites, further affects (mainly reduces) the transcription of certain housekeeping genes (especially carbon metabolism-related) that are actively transcribed in FL state | - |
| blr4773 | nwsA | T | N | 10 | m2 m4 m6 blr1499 blr2037 | Two-component hybrid sensor and regulator | Coordination between nitrogen fixation, nodulation and nitrogen assimilation. | [1-8] |
| blr0742 | ihfB | L | Y | 20 | m1 m7 m8 blr3780 | Integration host factor subunit beta | Influencing (mainly enhancing) gene expression | [9] |
| blr3780 | nadE | H | N | 8 | m3 m6 blr1499 blr0742 | NAD synthetase | - | - |
| blr1499 | exoN | M | Y | 11 | m2 m7 blr4773 blr3780 | UTP-glucose-1-phosphate uridylyltransferase | - | - |
| bll4944 | clpP | OU | N | 5 | m1 m3 m5 | ATP-dependent Clp protease proteolytic subunit | Hydrolysis of abnormal SNF associated proteins | [10] |
| blr0725 | ptsN | GT | Y | 8 | m2 m4 m6 | Nitrogen regulatory IIA protein | - | - |
| bll5385 | rplF | J | N | 6 | m3 m7 m8 | 50S ribosomal protein L6 | - | - |
| blr8147 | - | G | Y | 7 | m1 m2 bll7349 | PTS system transporter subunit IIA | - | - |
| blr2037 | nifA | KT | Y | 8 | m2 m6 blr4773 | Nif-specific regulatory protein | Coordination between nitrogen fixation, nodulation and nitrogen assimilation. | [1-8] |
| blr7448 | prsA | FE | N | 5 | m4 m8 | Ribose-phosphate pyrophosphokinase | - | - |


1. Grob, P. et al. A novel response-regulator is able to suppress the nodulation defect of a Bradyrhizobium japonicum nodW mutant. *Mol Gen Gene.* 1993, 241: 531-541
2. Loh, J and Stacey G. Nodulation Gene Regulation in Bradyrhizobium japonicum: a Unique Integration of Global Regulatory Circuits. *Appl Environ Microb*. 2003, 69: 10-17
3. Nellen-Anthamatten, D. et al. Bradyrhizobium japonicum FixK2, a crucial distributor in

the FixLJ-dependent regulatory cascade for control of genes inducible by low oxygen levels. *J. Bacteriol*. 1998, 180: 5251–5255.
4. Dixon R. and D. Kahn. Genetic regulation of biological nitrogen fixation. *Nat. Rev. Microbiol.* 2004. 2: 621–631
5. Mesa S. et al. Comprehensive Assessment of the Regulons Controlled by the FixLJ-FixK2-FixK1 Cascade in Bradyrhizobium japonicum. *J. Bacteriol*. 2008, 190: 6568–6579
6. Lindemann A, Moser A, Pessi G, Hauser F, Friberg M, Hennecke H, Fischer HM. New target genes controlled by the Bradyrhizobium japonicum two-component regulatory system RegSR. *J Bacteriol*. 2007, 189: 8928-8943
7. Martin G. B. et al. Role of the Bradyrhizobium japonicum ntrC Gene Product in Differential Regulation of the Glutamine Synthetase II Gene (glnII). *J. Bacteriol*. 1988, 170: 5452-5459
8. Franck W. L. et al. DNA Microarray-Based Identification of Genes Regulated by NtrC in Bradyrhizobium japonicum. *Appl Environ Microb*. 2015, 81: 5299 –5308.
9. Stonehouse E. et al. Integration Host Factor Positively Regulates Virulence Gene Expression in Vibrio cholerae. *J. Bacteriol*. 2008, 190: 4736–4748
10. Wang J. et al. The Structure of ClpP at 2.3 A Resolution Suggests a Model for ATP-Dependent Proteolysis. *Cell*. 1997, 91: 447–456

**Supplementary file.xlsx**

The reconstructed protein-protein interaction networks of *Bradyrhizobium diazoefficiens* USDA110 in different situations.
